# Multinucleated giant cells are hallmarks of ovarian aging with unique immune and degradation-associated molecular signatures

**DOI:** 10.1101/2024.12.03.626649

**Authors:** Aubrey Converse, Madeline J. Perry, Shweta S. Dipali, Jose V. V. Isola, Emmett B. Kelly, Joseph M. Varberg, Mary B. Zelinski, Jennifer L. Gerton, Michael B. Stout, Michele T. Pritchard, Francesca E. Duncan

## Abstract

The ovary is one of the first organs to exhibit signs of aging, characterized by reduced tissue function, chronic inflammation, and fibrosis. Multinucleated giant cells (MNGCs), formed by macrophage fusion, typically occur in chronic immune pathologies, including infectious and non-infectious granulomas and the foreign body response^1^, but are also observed in the aging ovary^2–4^. The function and consequence of ovarian MNGCs remain unknown as their biological activity is highly context-dependent, and their large size has limited their isolation and analysis through technologies such as single-cell RNA sequencing. In this study, we define ovarian MNGCs through a deep analysis of their presence across age and species using advanced imaging technologies as well as their unique transcriptome using laser capture microdissection. MNGCs form complex interconnected networks that increase with age in both mouse and nonhuman primate ovaries. MNGCs are characterized by high *Gpnmb* expression, a putative marker of ovarian and non-ovarian MNGCs^5,6^. Pathway analysis highlighted functions in apoptotic cell clearance, lipid metabolism, proteolysis, immune processes, and increased oxidative phosphorylation and antioxidant activity. Thus, MNGCs have signatures related to degradative processes, immune function, and high metabolic activity. These processes were enriched in MNGCs compared to primary ovarian macrophages, suggesting discrete functionality. MNGCs express CD4 and colocalize with T-cells, which were enriched in regions of MNGCs, indicative of a close interaction between these immune cell types. These findings implicate MNGCs in modulation of the ovarian immune landscape during aging given their high penetrance and unique molecular signature that supports degradative and immune functions.

## RESULTS AND DISCUSSION

### 3D networks of multinucleated giant cells are characteristic of the aging mammalian ovary

We undertook the first comprehensive assessment of ovarian multinucleated giant cells (MNGCs) to define their molecular signature and provide novel insight into their potential function and consequence in ovarian aging. To this end, we quantitatively assessed ovarian MNGCs across age in mouse and nonhuman primate (NHP) ovaries, imaged MNGCs in 3-dimensions using a tissue clearing approach, delineated the MNGC-enriched transcriptome using laser capture microdissection, and compared differentially expressed and unique genes between MNGCs and primary ovarian macrophages (Fig. 1A).

**Figure 1.**
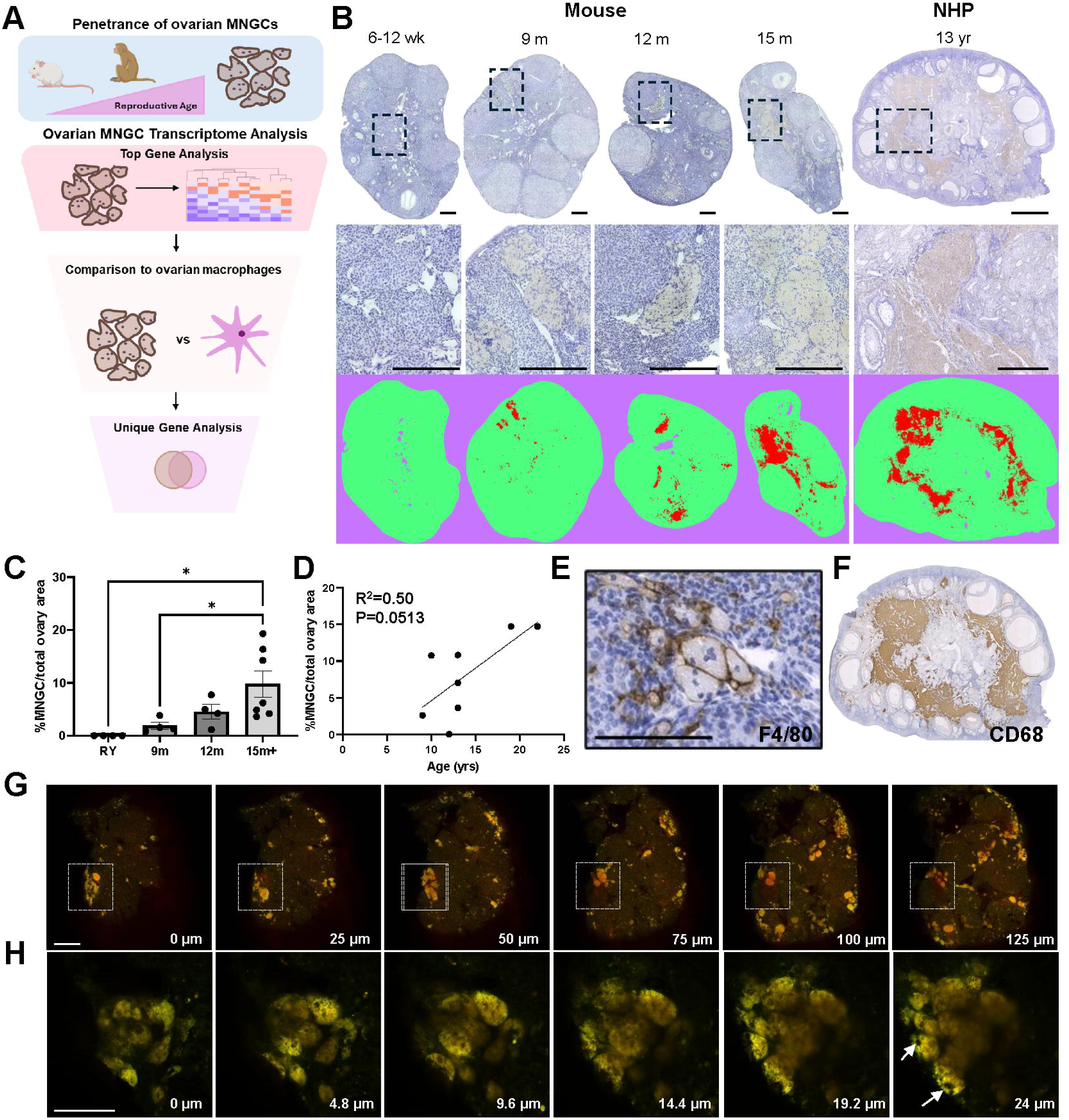
Multinucleated giant cells (MNGCs) are age-associated in mammalian ovaries. (A) Analysis workflow of the study. (B) Representative mouse and nonhuman primate (NHP) ovarian histological sections (top and middle panels) and Trainable Weka Segmentation (ImageJ) of MNGCs (bottom panel) across age. (C) Quantification of MNGC area in mouse ovaries across age. (D) Quantification of MNGC area in rhesus macaque ovaries across age. (E) F4/80 immunolabeling of mouse ovarian MNGCs. (F) CD68 immunolabeling of NHP MNGCs. Zstack of autofluorescent MNGCs in cleared mouse ovaries at macro (G) and micro (H) scales. Scale bars for B are=200 µm mouse top and middle, 2000 µm for NHP top panel, 650 µm for NHP middle panel. Scale bars=100 µm for E, 200 µm for G, and 50 µm for H. For C, data is presented as means ± SEM with 4-7 mice of each age. One-way ANOVA with Tukey’s multiple comparison post-test (*, p<0.05) was used for statistical analysis of C while a simple linear regression was used to determine the goodness to fit (R^2^) and the likelihood of the slope being non-zero (P) in D.

To characterize the penetrance of ovarian MNGCs across reproductive age, we quantified the percent MNGC area in ovarian tissue sections from mice and NHP across an aging spectrum. Ovarian MNGCs were absent in reproductively young (6-12wk) mouse ovaries but present in all ovaries from mice ≥ 9m of age (Fig. 1B & C). MNGCs occupied 1.9 ± 0.7, 4.5 ± 2.7, and 9.8 ± 6.6% of ovarian tissue sections at 9, 12, and 15m+, respectively. Ovarian MNGCs consistently increased between the 9 and 15+ month cohorts, and this was significant when comparing the young control and 9m groups with the 15+ month group. MNGCs were also present in NHP ovaries in animals aged 9-22 yrs, with an increasing trend with advanced age (Fig. 1B & D). Thus, the appearance of MNGCs is a hallmark of mammalian ovarian aging. We confirmed that ovarian MNGCs were, at least in part, derived from macrophages as they expressed F4/80 (mouse) and CD68 (NHP) (Fig. 1E-F), consistent with previous reports^2^. As ovarian MNGCs have only been visualized in tissue sections to date, it is unclear how they localize within the tissue and how vast individual MNGCs are in 3-dimensional (3D) space. Previous studies have demonstrated MNGCs are autofluorescence^2^ likely due to their high lipofuscin content^7^, so we utilized this autofluorescence to visualize MNGCs within cleared mouse ovarian tissue using multiphoton microscopy. Using this approach, we found that MNGCs formed large, interconnected networks within the ovary, with each cluster spanning hundreds of microns (Fig. 1G, SV1). On a microscale, we found that MNGCs were comprised of a complex autofluorescent cytoplasmic network containing numerous nuclei which was intercalated within non-autofluorescent tissue (Fig. 1H, SV2). To our knowledge, this is the first 3D mapping of native MNGCs within their resident tissue.

### LCM and RNA sequencing of ovarian multinucleated giant cells

Comprehensive transcriptomic analysis of ovarian and non-ovarian MNGCs has been limited in part because their large size excludes them from single-cell RNAseq preparations, and there is a lack of established primary MNGC isolation methods. Therefore, the field has heavily relied on *in vitro*-induced macrophage fusion models to assess the molecular characteristics and biological activity of MNGCs^8–10^. However, without direct comparison to *in vivo*-derived MNGCs, it remains unclear how well *in vitro* models reflect physiological counterparts. Although some studies have evaluated the transcriptome of native non-ovarian MNGCs^5,6^, it remains unclear how similar they are to ovarian MNGCs. We, therefore, utilized laser capture microdissection (LCM) to isolate native MNGCs from frozen mouse ovarian tissue (Fig 2A). MNGC-enriched regions were identified in tissue sections based on their distinct morphology and pigmented appearance in adjacent H&E-stained serial sections as well as their autofluorescence (Fig 2B).

**Figure 2.**
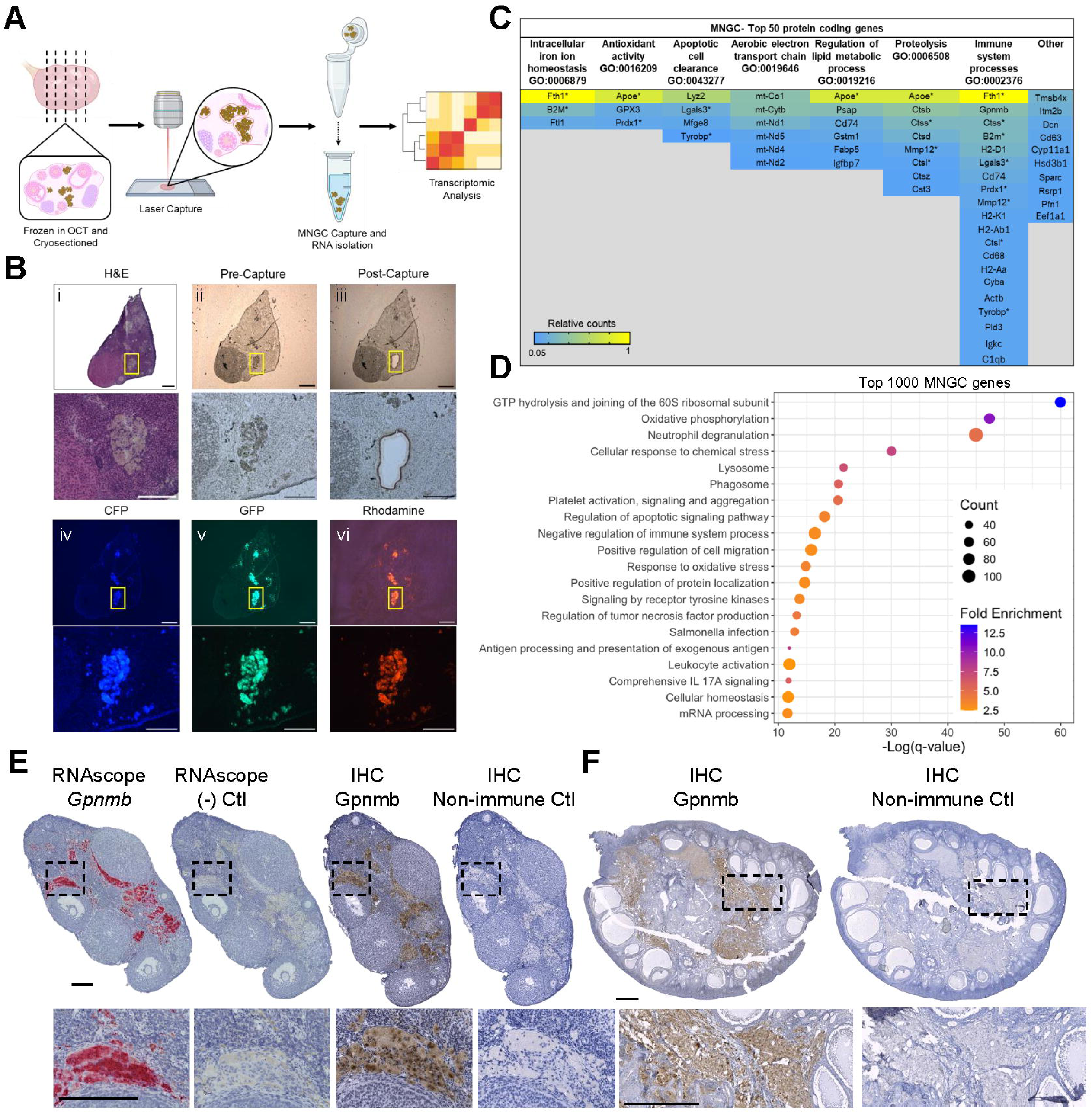
Transcriptomic analysis of ovarian MNGCs. (A) Schematic of laser capture microdissection (LCM) of ovarian MNGCs for transcriptomic analysis. (B) Representative images depicting MNGC identification by serial H&E-stained section (i) and autofluorescence (iv-vi), and capture for LCM (ii-iii). (C) Table of the top 50 MNGC expressed genes categorized by biological processes to which they relate. (D) Pathway analysis of top 1000 expressed MNGC genes. (E) Localization of *Gpnmb* RNA (RNAscope) and protein (IHC) in mouse (12m) ovarian sections. (F) Localization of GPNMB protein in rhesus macaque (13yr) ovarian tissue section. For C, (*) denotes genes that fall into more than one process. Scale bars=150 µm for B; 200 µm for E; 1000 µm for F. Transcriptomic analysis was performed on 4 biological replicates.

RNA sequencing revealed 9966 protein coding genes (≥10 counts) identified in all biological samples. We manually characterized the top 50 expressed genes into seven biological processes that were cross-referenced to annotated gene ontology (GO) terms (Fig. 2C). These included “intracellular iron ion homeostasis,” “antioxidant activity,” “apoptotic cell clearance,” “aerobic electron transport chain,” “regulation of lipid metabolic process,” “proteolysis,” and “immune system processes.” Of the top 50 genes, 80% fell into these processes, while the remainder were primarily involved in actin binding and matrix assembly (*Tmsb4x*, *Dcn*, *Sparc*, and *Pfn1*) or steroidogenesis (*Cyp11a1* and *Hsd3b1*). To further validate these processes, we performed unbiased pathway analysis of the top 1000 genes, which further highlighted immune, degradation, and oxidative phosphorylation processes (Fig. 2D). GO terms associated with degradation included “lysosome,” “phagosome,” and “regulation of apoptotic signaling pathway.” Immune function-related GO terms included “neutrophil degranulation,” “negative regulation of immune system processes,” “regulation of tumor necrosis factor production,” “*Salmonella* infection,” “antigen processing and presentation of exogenous antigen,” “leukocyte activation,” and “comprehensive IL17A signaling.” “Oxidative phosphorylation” and “response to oxidative stress” GO terms were also enriched. Thus, the biological processes manually annotated from the top 50 expressed genes were well-represented in an unbiased pathway analysis of the top 1000 genes.

From the top gene list, we selected *Gpnmb* for further characterization as it is only expressed in a small subset of ovarian macrophages^7^, is upregulated with age in the mouse proteome and transcriptome^11,12^, and is a putative marker of non-ovarian MNGCs^5,6^. RNA *in situ* hybridization and immunohistochemistry (IHC) demonstrated that, consistent with our transcriptomic data, *Gpnmb* transcripts localized within ovarian MNGCs and GPNMB protein was primarily localized to regions of MNGCs (Fig. 2E-F). These results indicate that MNGCs are the main cell type that expresses this marker and that the age-associated increase in ovarian GPNMB expression is likely due to the age-associated accumulation of MNGCs^11^. GPNMB is a trans-membrane glycoprotein implicated in processes such as immunosuppression in pro-inflammatory conditions^13,14^, modulation of ECM production and degradation^15–17^, and regulation of phagosome and lysosome function in debris clearance^18–21^. The protein is also upregulated in various cancers, autoimmune disorders, and senescent cells^20,22–28^. Interestingly, GPNMB is also expressed in osteoclasts where it promotes osteoclast precursor fusion and bone resorption activity^29^. GPNMB expression in various MNGC models, including the aging ovary, suggests that it likely has a specific role in MNGCs, potentially related to immune signaling, degradation, and fusion.

### Ovarian MNGCs are molecularly distinct from ovarian macrophages

To identify potential novel functions of MNGCs distinct from other ovarian macrophage populations, we compared the transcriptomic signatures of MNGCs and ovarian macrophages isolated from reproductively young mice. Young ovarian macrophages were utilized as a control to minimize any confounding influence of age-associated changes on macrophage populations and transcriptomic profiles^30–32^. Ovarian macrophages were isolated by enzymatic digestion of ovaries, filtering to obtain a single-cell suspension, and immunomagnetic pull-down of F4/80+ cells (Fig. 3A). A subset of cells isolated by immunomagnetic sorting were utilized to confirm their identity based on phagocytic activity (Fig. 3B). Initial comparison of the transcriptomic profiles of ovarian MNGCs and ovarian macrophages revealed discrete separation between these populations in a principal component analysis (PCA; Fig. 3C). Next, the expression of macrophage markers was compared between macrophage populations and stroma samples that were collected by LCM, which served as a non-macrophage enriched population (Fig 3D). Both macrophage populations had higher expression of the pan-macrophage marker *Adgre1* (F4/80) than stroma samples. Expression of macrophage-associated genes such as *CD14*, *Nos2*, and *Tlr2* were significantly higher in macrophages than MNGCs or stroma. In contrast, other markers, including *Cd68*, *Lgals3, and Gpnmb,* were significantly higher in MNGCs compared to macrophages or stroma. Overall, these data confirm the macrophage identity of the F4/80 immuno-isolated cells and MNGCs.

**Figure 3.**
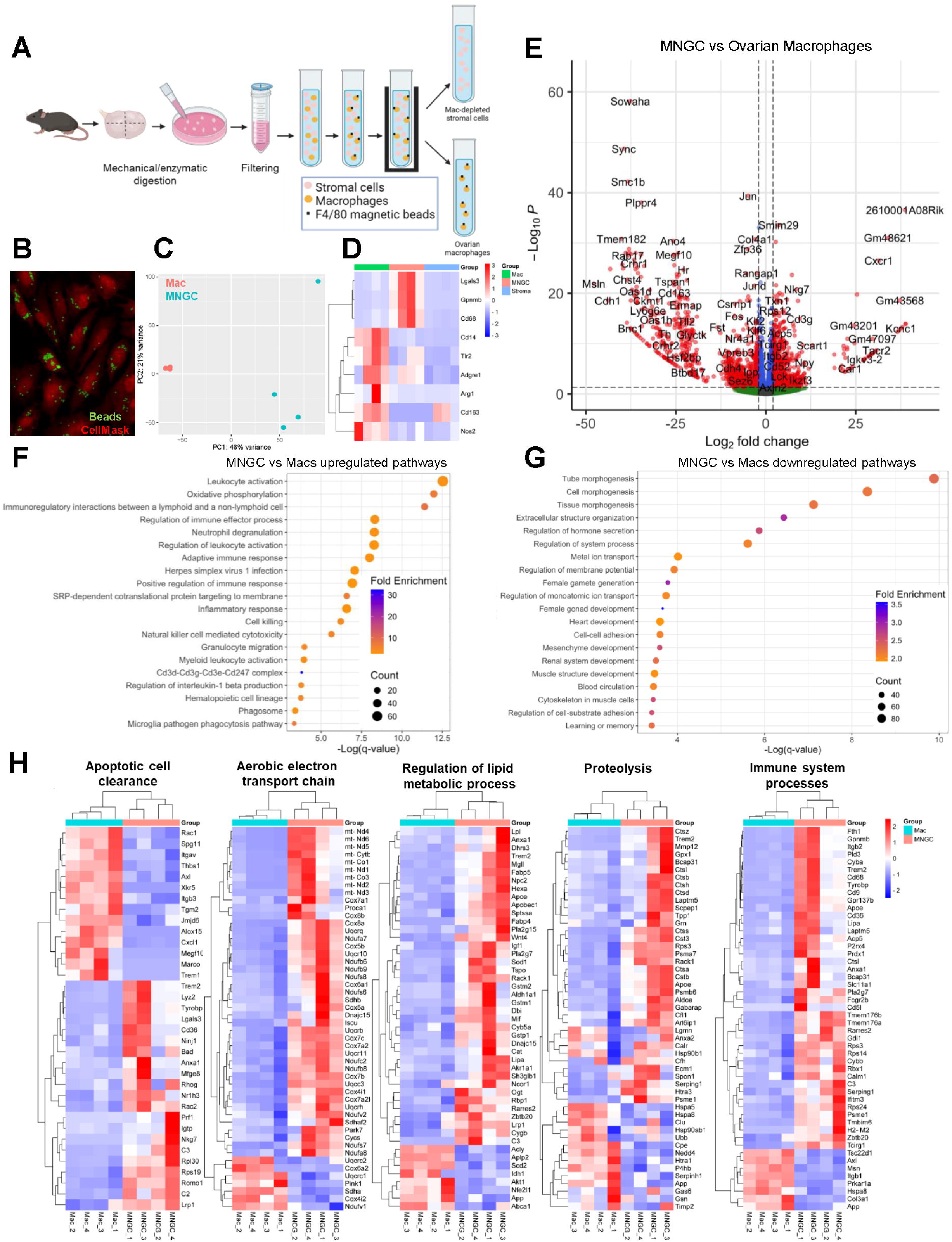
Comparison of ovarian MNGCs and primary ovarian macrophages. (A) Schematic of isolation procedure for ovarian macrophages by immunomagnetic pull-down. (B) Confirmation of macrophage identity of F4/80-pull down cells by phagocytic capacity. (C) Principal component analysis (PCA) of primary ovarian macrophage (Mac) and MNGC transcriptomes. (D) Expression profiles of macrophage marker genes in primary macrophages, MNGCs, and non-MNGC stromal samples. (E) Volcano plot of differentially expressed genes (DEGs) between MNGCs and primary ovarian macrophages (p<0.05 and Log_2_FC≥2). (F) Pathway analysis of upregulated DEGs in MNGCs compared to primary macrophages. (G) Pathway analysis of downregulated DEGs in MNGCs compared to primary macrophages. (H) Heat maps of MNGC vs primary macrophage DEGs for select biological processes. Transcriptomic analysis was performed on 4 biological replicates for each sample type.

Transcriptomic analysis identified 5436 differentially expressed genes (DEGs, p ≤ 0.05) between ovarian macrophages and MNGCs, of which 1239 were upregulated and 1390 were downregulated at a log_2_ fold-change (FC) ≥ 2 (Fig. 3E). GO analysis of genes upregulated in MNGCs indicated substantial enrichment for immune-related processes such as “leukocyte activation,” “adaptive immune response,” and “inflammatory response”, among others (Fig. 3F). “Oxidative phosphorylation” and “phagosome” GO terms were also enriched. Processes identified from DEGs that were downregulated in MNGCs compared to macrophages were enriched in GO terms relating to morphogenesis and development and were driven by genes related to development such as *Notch1*, *Foxc1* and *Wnt7b* (Fig. 3G). These results may be indicative of decreased phenotypic plasticity of MNGCs compared to macrophages.

Heatmaps displaying relative expression of genes within biological processes supported by biased and non-biased pathway analysis further highlighted distinct differences in functionality between MNGCs and ovarian macrophages (Fig. 3H). Biological pathways enriched in MNGCs compared to macrophages were aligned with those identified by top expressed MNGC genes (Fig. 2D), indicating that these findings are not reliant on the comparison to or the expression profile of ovarian macrophages. Furthermore, these pathways agree with known functions of non-ovarian MNGCs. In other tissues, MNGCs play major roles in phagocytosis, degradation, and immune signaling, though their specific roles differ depending on their context^1^. Recent transcriptomic analyses indicated foreign body giant cells (FBGCs) have increased expression of genes related to lipid and glucose metabolism and extracellular matrix degradation compared to other immune and non-immune cell types^5^, whereas MNGCs of sarcoidosis are enriched for lysosomal and phagosome processes^6^. In *in vitro*-derived MNGCs, processes such as fatty acid and cholesterol metabolism, lysosome, and iron transport are upregulated during multinucleation^9,33^. The enrichment of pathways involved in degradation in ovarian and non-ovarian MNGCs indicate this is likely a conserved function of all MNGCs. Furthermore, the appearance of MNGCs in the aging ovary may indicate that aging results in disruption of the debris clearance mechanisms in the ovary, a highly dynamic organ that undergoes cyclic cell proliferation and death each estrous/menstrual cycle. Increased degradative function likely requires increased energy production, as is the case in osteoclasts in which increased ATP-generation by oxidative phosphorylation is vital for survival and bone degradation activity^34,35^. Increased oxidative phosphorylation likely also explains the upregulation of antioxidant genes as a reflection of increased ROS production.

### Ovarian MNGCs express CD4 and colocalize with CD3+ cells

To assess unique features of the ovarian MNGC and macrophage transcriptomes, we compared all protein coding genes between these groups. While 9827 genes were shared by both macrophage populations, ovarian macrophages expressed 3311 genes that were absent from the MNGC group, whereas MNGCs expressed 137 unique genes (Fig. 4A). GO analysis of the genes unique to ovarian macrophages indicated enrichment for pathways involved in cell cycle, development and morphogenesis, and cell migration processes (Fig. 4B). Pathway analysis of genes unique to MNGCs revealed enrichment for “Cd3d-Cd3g-Cd3e-Cd247 complex” and “cytokine-cytokine receptor interaction” (Fig. 4C). Upon examination of the 15 unique MNGC genes with the highest expression, all CD3 complex components were present as well as genes involved in modulation of T-cell activity such as *Nkg7*, *Sh2d2a*, and *Ctsw* (Fig. 4D). As CD3 is a T-cell co-receptor typically only expressed in T-cells, we examined expression of various T-cell markers between the MNGC, ovarian macrophages, and stroma samples (Fig. 4E). MNGC samples had significantly higher expression of *Cd3d* and *Cd3e* and *Trbc2* compared to stroma and macrophage samples, with increases ranging between 8 to 633-fold enrichment. The other CD3 complex member *Cd3g*, the cytotoxic T-cell marker *Cd8a*, the helper T-cell marker *Cd4*, and other TCR constant chain genes showed trends in increased expression in MNGC samples compared to normal macrophage and stromal samples.

**Figure 4.**
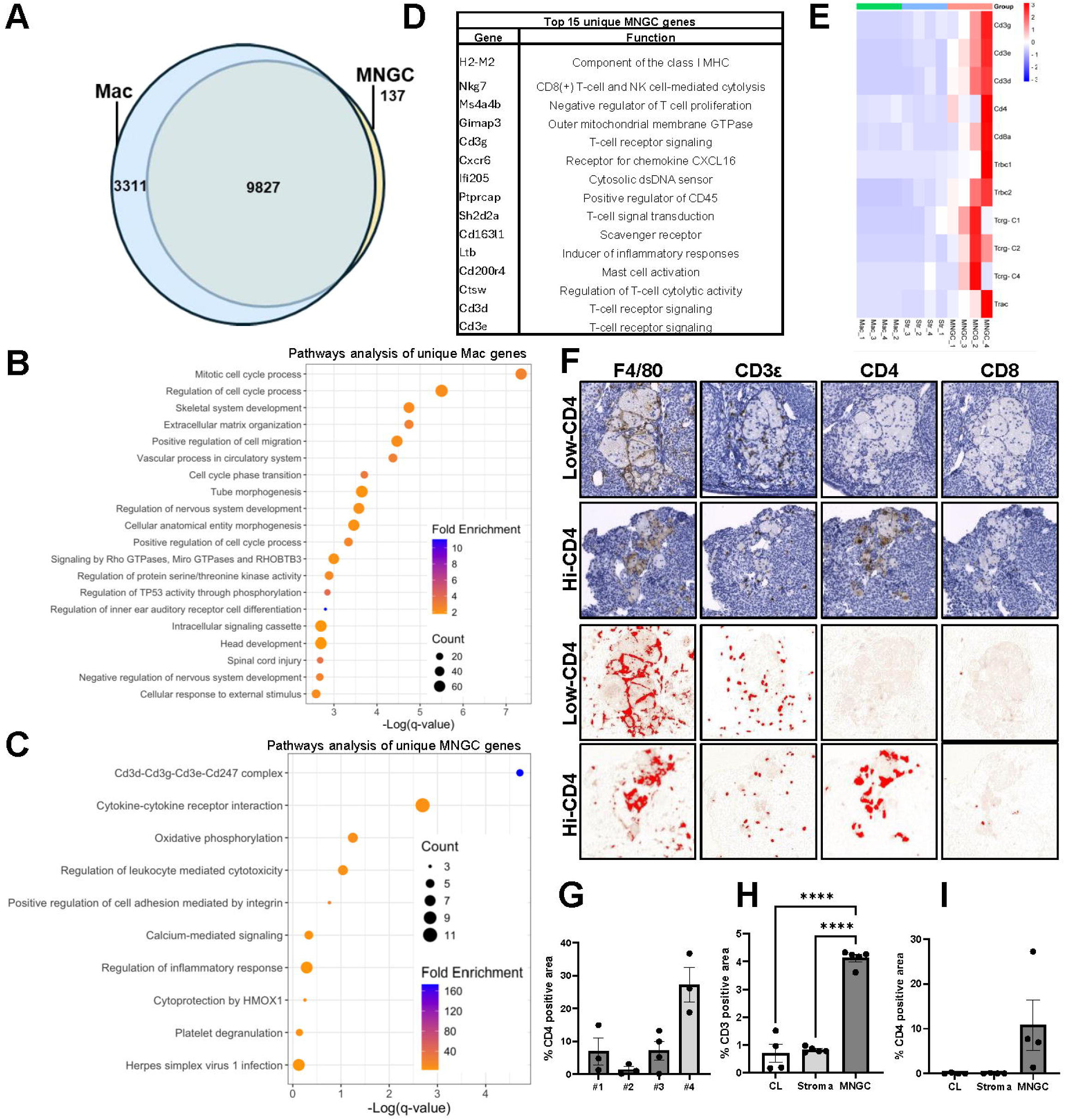
Assessment of gene signatures unique to MNGCs. (A) Venn diagram of protein coding genes (≥10 counts in all samples) in primary macrophages and MNGCs. (B) Pathway analysis of genes unique to ovarian macrophages from young mice. (C) Pathway analysis of genes unique to ovarian MNGCs. (D) Top 15 expressed unique MNGCs genes. (E) Heat map of expression profiles of T-cell marker genes. (F) Immunohistochemical analysis of macrophage (F4/80) and T-cell (CD3ε, CD4, CD8a) maker protein expression in ovarian MNGCs. (G) CD4 positive area of MNGCs from 3-4 individual MNGCs in four biological samples (#1-4). Quantification of CD3ε (H) and CD4 (I) positivity of MNGCs, stroma, and corpora lutea. For G-I, data represents means ± SEM. For H and I, data were analyzed by one-way ANOVA with Tukey’s multiple comparison post-test (****, p<0.0001). All transcriptomic and histological analyses were performed on 4 biological replicates.

To further understand why T-cell specific markers were detected in the transcriptome of MNGCs, we immunolocalized CD3ε, CD4, and CD8 in ovaries from 12m old animals via IHC (Fig 4F). While all MNGCs examined stained positive for F4/80, confirming their identity as macrophages, CD3 positive cells were consistently found to be intercalated within regions of MNGCs. These cells were observed to be outside of shared cytoplasmic regions, indicating that they likely are not fused or within MNGCs, but rather cells in between the 3D interconnected cytoplasmic network (Fig. 1G-H). Serial sections in which CD4 and CD8 were immunolocalized did not demonstrate a similar punctate staining pattern as CD3ε, indicating the majority of the MNGC-associated CD3+ cells are likely CD4 and CD8 negative T-cells (double negative T-cells; dnT). dnT cells increase with age in the ovary and can originate from thymocyte double negative precursors or downregulation of CD4 or CD8 in peripheral T-cells^7,32,36–38^. While dnTs have both innate and adaptive immune capabilities^39^, their consequence in the aged ovary is unknown.

Although CD3+ cells closely associate with MNGCs, a portion of the F4/80+ regions of MNGCs also exhibited CD4 immunoreactivity, indicating that MNGCs can express this T-cell co-receptor (Fig. 4F). CD4 positive MNGC regions were highly variable between animals and between different MNGCs within the same animal (Fig. 4G), with CD4 positivity of MNGCs ranging from 0.1% to 36.7%. To determine if MNGCs showed an enrichment of T-cell markers compared to other ovarian structures, the positive signals for CD3ε and CD4 were quantified in ROIs of MNGCs, corpora lutea, and non-MNGC regions of stroma. CD3ε immunoreactivity was significantly enriched in MNGCs compared to other ovarian structures, with a 5.0 and 5.9-fold enrichment compared to regions of stroma and corpora lutea, respectively (Fig. 4H; Supplemental Fig. 1A). MNGCs demonstrated a trend in increased CD4 immunoreactivity compared to other ovarian compartments (Fig. 4I; Supplemental Fig. 1B). Thus, MNGCs represent regions in which T-cells and T-cell markers display enrichment compared to the broader ovarian microenvironment.

To further explore cell-to-cell interactions between MNGCs and T-cells, we utilized a previously established single-cell RNAseq dataset of mouse ovarian aging^7^. While large MNGCs are excluded from single-cell data sets, we routinely observed autofluorescent MNGCs in aged ovaries that were <40 µm (Supplemental Fig. 2A) and that would be capable of passing through single-cell flow cells. As *Gpnmb* expression is a conserved marker of different MNGC populations^5,6^, we utilized *Gpnmb* positivity as an indicator of MNGC-like cells. *Gpnmb* was only expressed by a small fraction of the macrophage subcluster (Supplemental Fig. 2B-C) and displayed an increasing trend with age (Supplemental Fig. 2D), supporting that MNGC-like cells are represented in this fraction. CellChat^40^ analysis of interactions between *Gpnmb*(+) macrophages and immune subclusters indicated high levels of self-signaling, moderate interactions with Type I and Type II lymphoid cells, and lower levels of interaction with CD4+ cells and CD45-negative immune like cells (subcluster A) (Supplemental Fig. 2E). MNGC-like cell interaction with both lymphoid populations showed higher weights of outgoing signals compared to incoming signals, suggesting that MNGCs may modulate lymphoid cell behavior to a greater degree than lymphocytes modulate MNGC activity. Of note, many of the cells included in both lymphoid subclusters contained CD4-CD8-cells, further supporting the observed association of dnT cells with MNGCs in the current study.

The close association of T-cells with MNGCs in this system shares similarities with giant cells of granulomas, in which various immune cell types are present. T-cells are implicated in granuloma formation with T-cell deficient animal models exhibiting abrogated granuloma formation^41–43^. Furthermore, cytokines produced by T-cells such as RANKL and IL-4 can induce macrophage fusion^9^. However, the role of T-cells in MNGC formation does not appear to be critical as T-cell deficiency has little effect on foreign giant cell formation *in vivo*^44^. T-cell recruitment to MNGC-rich areas by MNGC chemokine secretion and their potential to act as antigen presenting cells may also drive the close association, as these immunoregulatory functions are well characterized in osteoclasts^1,45–47^. An immunoregulatory role of ovarian MNGCs is supported by their expression of CD4 and their enrichment for genes involved in adaptive immune signaling. Additionally, the tight co-localization of CD3+ cells and MNGCs suggests that MNGCs may be a product or driver of the age-associated T-cell infiltration to the ovary^7,32^. In either case, while the infiltrating CD3+ cell signature is present in our MNGC transcriptomic data, it reveals the comprehensive signal of this broader age-associated immune cell complex. Additionally, pathway analysis after exclusion of genes with counts below that of the highest expressed T-cell specific gene *Nkg7*, further confirm degradation and unique immune functions can be attributed directly to MNGCs (Supplemental Tables 1 and 2).

In this study, we comprehensively characterized and defined the molecular signature of age-associated ovarian MNGCs, supporting their role in degradation and immune signaling within the aging ovary (Graphical Abstract). Ovarian MNGCs share similarities with non-ovarian MNGCs, such as granuloma, foreign body response, and osteoclast giant cells, but also exhibit unique immune characteristics. Their unique immune signature and the close association with T-cell populations indicates ovarian MNGCs likely play a role in the robust changes to the ovarian immune landscape with aging^2,7,30–32^.

## METHODS

### Animals

All animals used in this study were C57BL/6J (transcriptomic analyses) and C57BL/6Hsd (histological analyses) and were acquired from the JAX aged colony (Jackson Laboratory, Bar Harbor, ME) or Envigo (Indianapolis, IN), respectively. Animals were maintained in accordance with the National Institutes of Health’s guidelines and housed in Northwestern University’s Center for Comparative Medicine barrier facility under constant light (14h light/10 hr dark), humidity, and temperature control, and offered food and water *ad libitum*. All animal experiments were approved by Northwestern University’s Institutional Animal Care and Use Committee.

### Preparation of samples for histological analyses

For mouse samples, ovaries were collected from animals aged 8-10 wk, 9-10m (9m), 12-13m (12m), 15-19m (15m+), fixed in Modified Davidson’s (Electron Microscopy Sciences, Hatfield, PA, USA) and processed for histological analysis as previously described^2^. All mice were sampled while in diestrus to account for potential effects of ovarian cyclicity. Rhesus macaque (Macaca mulatta; nonhuman primate (NHP)) samples were generously provided by Dr. Mary Zelinski. NHP ovaries were fixed in 4% paraformaldehyde (Sigma-Aldrich, St. Louis, MO) for 24 hours at 4°C, equilibrated in 4% sucrose (Sigma-Aldrich) for 24 hr at 4°C, then stored in 70% ethanol prior to dehydration, embedding in paraffin, and sectioning at 5 µm.

### MNGC quantification

Slides were stained with Harris hematoxylin (EK industries Inc., Joliet, IL) and imaged with an EVOS FL Auto Imaging system (Thermo Fisher Scientific, Waltham, MA) at 20X. To quantify the amount of ovarian area taken up by MNGCs, the Trainable Weka Segmentation plugin in Fiji (NIH) was used to train the model on regions of non-ovary, non-MNGC ovary, and MNGCs. For mouse samples, training and quantification was performed on 3 sections from different regions of the ovary that were at least 300 µm apart, with the average MNGC area per mouse reported. For NHP samples, one midsection per animal was assessed at each age for MNGC content.

### Ovarian clearing and multiphoton and confocal microscopy

Optical clearing was performed on 12m old C57BL/6Hsd ovaries using a MACS clearing kit (Miltenyi Biotec, Gaithersburg, MD) following the manufacturer’s protocol for clearing of mouse spleen, kidney, and liver samples. A multiphoton microscope with a Cousa air objective that allows for an ultra-long working distance was used^48^. Multiphoton images were acquired with a Nikon A1R MP system on a Nikon NiE upright stand equipped with gallium arsenide phosphide (GaAsP) detectors. A Coherent Chameleon Vision II laser was tuned to 850. The objective used was the 10x 0.5NA Cousa objective from Pacific Optica^48^ (Santa Barbara, CA). Zstack images were acquired every 0.8 µm (Fig. 1H) or 2.5 µm (Fig. 1G). For scanning laser confocal microscopy (Supplemental Fig. 2A), a Leica TCS Sp5 confocal system with 40X oil objective (Leica Microsystems, Deerfield, IL) was utalized.

### Laser capture microdissection of MNGCs

Ovaries from 18-20m old C57BL/6J mice were dissected and embedded in optimal cutting temperature (OCT) compound under RNAase-free conditions and stored at −80°C. Fresh-frozen ovarian tissue was cryosectioned at 10 µm, adhered to nuclease-free PEN membrane slides (Carl Zeiss Microscopy, Jena, Germany), and stored at −80°C. Tissue sections were fixed in 70% molecular grade ethanol for 3 minutes, transferred to RNase-free water to remove OCT, and then dehydrated by incubating in 70% and then 100% ethanol for 3 min each. Following dehydration, the slide was air dried and immediately used for laser capture microdissection (LCM) of MNGCs or non-MNGC stroma regions on a Zeiss PALM Laser Capture Microdissection System (Carl Zeiss Microscopy) using PALM RoboSoftware (Carl Zeiss Microscopy). All cuts were made while on a 20x objective, with a cut energy ranging between 48-54 and cut focus ranging between 62-69, an LPC energy of 25, and LPC focus of −3. Cuts were outlined using the RoboLPC setting and free-hand outlining tool. Once the region of interest was defined, tissue was captured into an AdhesiveCap 500 (Carl Zeiss Microscopy). Collection ceased after 40 min or when 250 µm^3^ of tissue had been collected. Collection tubes were snap frozen on dry ice and stored at −80°C.

### Ovarian macrophage isolation

Ovaries from 6-12wk C57BL/6J mice were used to eliminate the potential inclusion of smaller MNGCs or MNGC precursors, as well as negate any age-related effects on macrophage phenotype. Ovaries were cut into eighths and digested in Leibowitz L15 media containing 1% fetal bovine serum (FBS; Gibco, Waltham, MA), 0.5% penicillin-streptomycin, 0.4 mg/ml Collagenase IV (205 units/mg; Gibco), 40 µg/ml Liberase DH (Sigma Aldrich), and 0.2 mg/ml DNase I (Sigma Aldrich) for 30 min. Tissue was pipetted to mechanically disrupt at 0, 15, and 30 min during digestion. After digestion, sterile FBS was added to the cells at a 10% volume, cells were filtered through a 40 µm cell strainer, centrifuged at 300 x g for 5 min, then resuspended in 1 ml EasySep buffer (STEMCELL Technologies Inc., Cambridge, MA). Positive selection of macrophages was performed utilizing an EasySep Mouse F4/80 Positive Selection Kit (STEMCELL Technologies Inc.) following the manufacturer’s protocol. Immunoselected cells were immediately processed for RNA isolation or plated in Dulbecco’s Modified Eagle Medium (DMEM) with 10% FBS and 1% penicillin-streptomycin for subsequent assessment of functional activity.

### Phagocytosis assay

Cells isolated with immunomagnetic pull-down were plated on collagen (50 µg/ml; Corning, Corning, NY) coated coverslips overnight, incubated with fluorescent latex beads (1.0 µm; Sigma Aldrich) for 3 hrs, then washed extensively with PBS and fixed for 15 min with 4% paraformaldehyde. After three washes with PBS, coverslips were incubated with 1X HCS CellMask orange stain (Thermo Fisher Scientific) for 1hr, washed 3x with PBS, and mounted with Vectashield Plus Antifade Mounting Medium with DAPI (Vector Laboratories, Burlingame, CA). Cells were imaged with a Leica TCS Sp5 confocal system using 40X objective (Leica Microsystems, Deerfield, IL).

### RNA isolation and quality control

Total RNA was isolated from LCM-collected samples and primary macrophages using a RNeasy Micro Kit (Qiagen, Germantown, MD) following the manufacturer’s protocol for purification of total RNA from microdissection cryosections with on-column DNase digestion. For LCM-collected samples, 350 µl of RLT buffer containing β-Mercaptoethanol was added to the AdhesiveCap tube, vortexed for 30s, stored upside down for 30 min at room temperature, then centrifuged at 13,400 x g for 5 min. The samples were then moved to 1.5 ml RNase-free centrifuge tubes and immediately processed following the manufacturer’s protocol. All samples were eluted into 14 µl nuclease-free water and stored at −80°C. RNA integrity was assessed using a 2100 Bioanalyzer System (Agilent Technologies, Inc., Santa Clara, CA) with a Bioanalyzer RNA Pico chip (Agilent Technologies, Inc.). For primary macrophages, RIN values were between 8 and 9, while for LCM-collected MNGC samples RIN values ranged from 6.9-8.2.

### Library generation, bulk RNA sequencing, and bioinformatics analysis

Cell content was used as direct input for full length cDNA synthesis using SMART-Seq v4 Ultra Low Input RNA Kit for Sequencing (Takara Bio, San Jose, CA) according to manufacturer’s protocol. The resulting cDNA quantity and quality were measured by Qubit DNA HS assay and Agilent Bioanalyzer DNA HS chip, respectively. Only samples with high quality cDNA (average size above 800bp) were used to generate sequencing libraries. 200 ng of cDNA was used as input for library preparation using Illumina Nextera XT DNA library preparation kit (Illumina, Inc., San Diego, CA) according to manufacturer’s protocol. Multiplexed libraries from all samples (LCM collected and primary macrophages) were sequenced within a single flow cell on Novaseq X Plus at the Northwestern University NUSeq core facility. The read quality, in FASTQ format, was evaluated using FastQC. Reads were trimmed to remove Illumina adapters from the 3’ ends using cutadapt^49^. Trimmed reads were aligned to the *Mus musculus* genome (mm10) using STAR^50^. (http://useast.ensembl.org/index.html). Normalization was performed using DESeq2^51^. Differential expression analysis was performed using DESeq2 and the contrast argument of the results function was utilized to compare MNGCs to primary ovarian macrophages. Data were rlog transformed and PCA plots were generated using the plotPCA function. Volcano plots were generated using the EnhancedVolcano package in R and differentially expressed genes with adjusted p-values <0.05 and absolute log_2_ fold-change ≥2 were considered significant. All gene ontology analyses were performed using Metascape^52^ with top expressed genes, unique genes, or differentially expressed genes with adjusted p-values <0.05 and absolute log_2_ fold-change ≥2. Dot plots were created using the ggplot2 package in R to visualize gene ontology pathways. Heatmaps and Venn diagrams were generated using SRPlot and VennDetail Shiny, respectively. For Venn diagrams, only protein coding genes with normalized counts ≥10 for each biological sample were included. Bioinformatic data utilized from Isola et al. 2024 was generated and processed as described in the primary publication.

### RNAscope in situ hybridization and immunohistochemistry

*Gpnmb* transcripts were visualized on ovarian tissue sections using a RNAscope 2.5 HD Assay-RED kit (Advanced Cell Diagnostics, Newark, CA) following the manufacturer’s protocol. For immunohistochemistry, tissue sections were deparaffinized in Citrosolv, rehydrated in graded ethanol dilutions (100, 95, 85, 70, and 50%) and then washed in deionized water. Antigen retrieval was carried out by incubation in 1X Reveal Decloaker (Biocare Medical, Concord, CA) in a steamer for 30 min. Endogenous peroxide activity was blocked by incubation with 3% hydrogen peroxide for 15 min, and endogenous avidin and biotin was blocked using and avidin/biotin blocking kit (Vector Laboratories). Slides were blocked in 3% bovine serum albumin (BSA, Sigma-Aldrich) and 10% goat serum (Vector Laboratories) for 1 hr at room temp, prior to overnight incubation with primary antibodies for F4/80 (1:50), GPNMB (1:500), CD3ε (1:400), CD4 (1:100), or CD8 (1:50) diluted in 3% BSA at 4°C. Protein concentration-matched non-immune rabbit or rat IgG was used for negative controls. After primary antibody incubation, the sections were washed in TBS with 0.1% Tween 20 (Sigma-Aldrich) and incubated with a biotinylated anti-rabbit (1:200) or anti-rat (1:100) secondary antibody (Vector Laboratories) for 2 hr at room temperature. Signal amplification was performed with a Vectastain Elite ABC Kit (Vector Laboratories) and detection was performed with a 3,3’-diaminobenzidine (DAB) with the DAB Peroxidase (HRP) Substrate Kit (Vector Laboratories). Tissue sections were counterstained with Harris hematoxylin, dehydrated and mounted with Cytoseal XYL. Sections were imaged with an EVOS M7000 (Thermo Fisher Scientific) at 20X and percent positive CD4 or CD3ε area was determined as DAB positive area over the total area of an ROI that included area of MNGCs, CLs, or stroma. Two to three areas for each feature were assessed for each section, and sections from 4 animals were assessed for analyses.

### Statistical analysis

Statistical analysis was performed using GraphPad Prism 10.2 Software (Boston, MA) or R (RStudio, version 4.3.0). Statistical significance was determined by one-way ANOVA with post hoc Tukey’s multiple comparison test or simple linear regression with *P* values <0.05 considered significant. All data are expressed as the mean ± SEM and analysis was performed with samples from a minimum of four mice.

## ACKNOWLEDGEMNTS

This work was supported in part by the Global Consortium for Reproductive Longevity & Equality grants GCRLE-1223 (A.C. & M.T.P.) and GCRLE-4501 (M.B.S.), the National Institutes of Health (NIH) grants R01HD105752 (F.E.D. & J.L.G), AG069742 (M.B.S.), P51 OD011092 (DPCPS, ORIP, to the Oregon National Primate Research Center), and U01 AG021382 (M.B.Z.), and Northwestern University Startup funds to F.E.D. We also acknowledge Northwestern University’s NUseq and Center for Advanced Microscopy cores for help with RNAseq experiments and multiphoton microscopy imaging, respectively. We thank the Biology of Aging (BOA) course at the Marine Biological Laboratory for allowing us to pilot experimental designs, assays, and reagents, as well as BOA students Ziying Xu and Diego Acuna for their assistance with preliminary experiments. We acknowledge the Gerton Lab at the Stowers Institute (Kansas City, MO) and the Immunology Group at the University of Kansas Medical Center (Kansas City, KS) for their valuable input which helped guide this study. Multiphoton microscopy was performed at the Northwestern University Center for Advanced Microscopy (RRID: SCR_020996) generously supported by CCSG P30 CA060553 awarded to the Robert H Lurie Comprehensive Cancer Center using the Nikon A1RMP purchased with the support of NIH 1S10OD016342-01.

## AUTHOR CONTRIBUTIONS

Conceptualization, A.C., M.T.P., F.E.D.; Methodology, A.C., M.T.P., F.E.D.; Formal Analysis, S.S.D., J.V.V.I., E.K.; Investigation, A.C., M.J.P.; Writing – Original Draft, A.C.; Writing – Review & Editing, A.C., M.T.P., F.E.D., Funding Acquisition, A.C., M.T.P., F.E.D., M.B.S, M.B.Z; Resources, M.B.S., M.B.Z.; Visualization, M.J.P.; Provided Expertise, J.V.V.I., J.M.V., J.L.G., M.B.Z., M.B.S.; Supervision A.C.

## RESOURCE AVAILABILITY

### Lead Contact

Requests for further information and resources should be directed to and will be fulfilled by the lead contact, Aubrey Converse.

### Materials availability

This study did not generate new unique reagents.

### Data and code availability

The RNA sequencing data has been deposited GEO and made available upon acceptance.

This study did not generate any unique code.

Any additional information required to reanalyze the data reported in this work paper is available from the lead contact upon request.

