## Supplemental figures and tables for "Multinucleated giant cells are hallmarks of ovarian aging with unique immune and degradation-associated molecular signatures"

### Slide 1
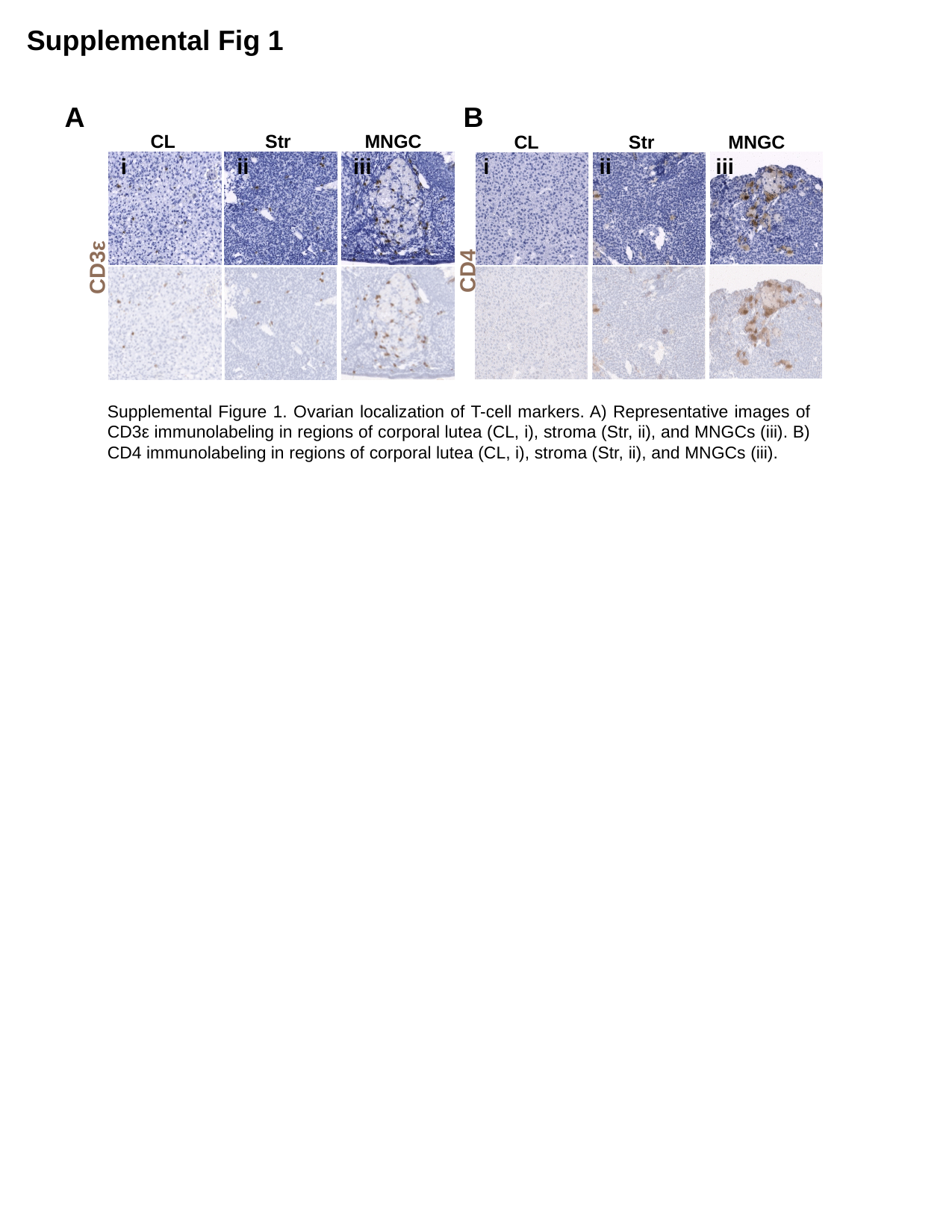

Supplemental Fig 1
A
B
CL
Str
MNGC
CL
Str
MNGC
i
ii
iii
i
ii
iii
CD3ε
CD4
Supplemental Figure 1. Ovarian localization of T-cell markers. A) Representative images of CD3ε immunolabeling in regions of corporal lutea (CL, i), stroma (Str, ii), and MNGCs (iii). B) CD4 immunolabeling in regions of corporal lutea (CL, i), stroma (Str, ii), and MNGCs (iii).

### Slide 2
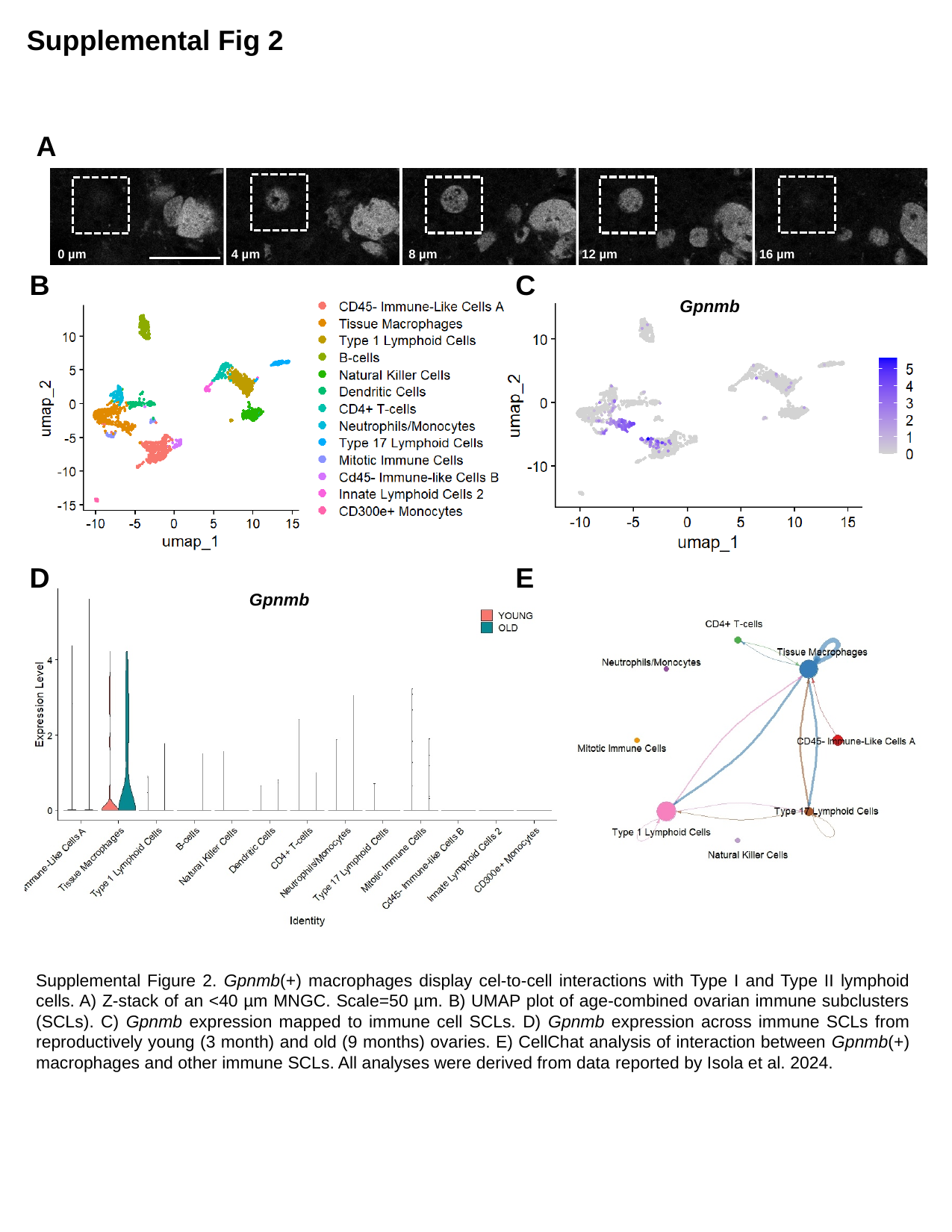

Supplemental Fig 2
A
0 µm
4 µm
8 µm
12 µm
16 µm
B
C
Gpnmb
D
E
Gpnmb
Supplemental Figure 2. Gpnmb(+) macrophages display cel-to-cell interactions with Type I and Type II lymphoid cells. A) Z-stack of an <40 µm MNGC. Scale=50 µm. B) UMAP plot of age-combined ovarian immune subclusters (SCLs). C) Gpnmb expression mapped to immune cell SCLs. D) Gpnmb expression across immune SCLs from reproductively young (3 month) and old (9 months) ovaries. E) CellChat analysis of interaction between Gpnmb(+) macrophages and other immune SCLs. All analyses were derived from data reported by Isola et al. 2024.

### Slide 3
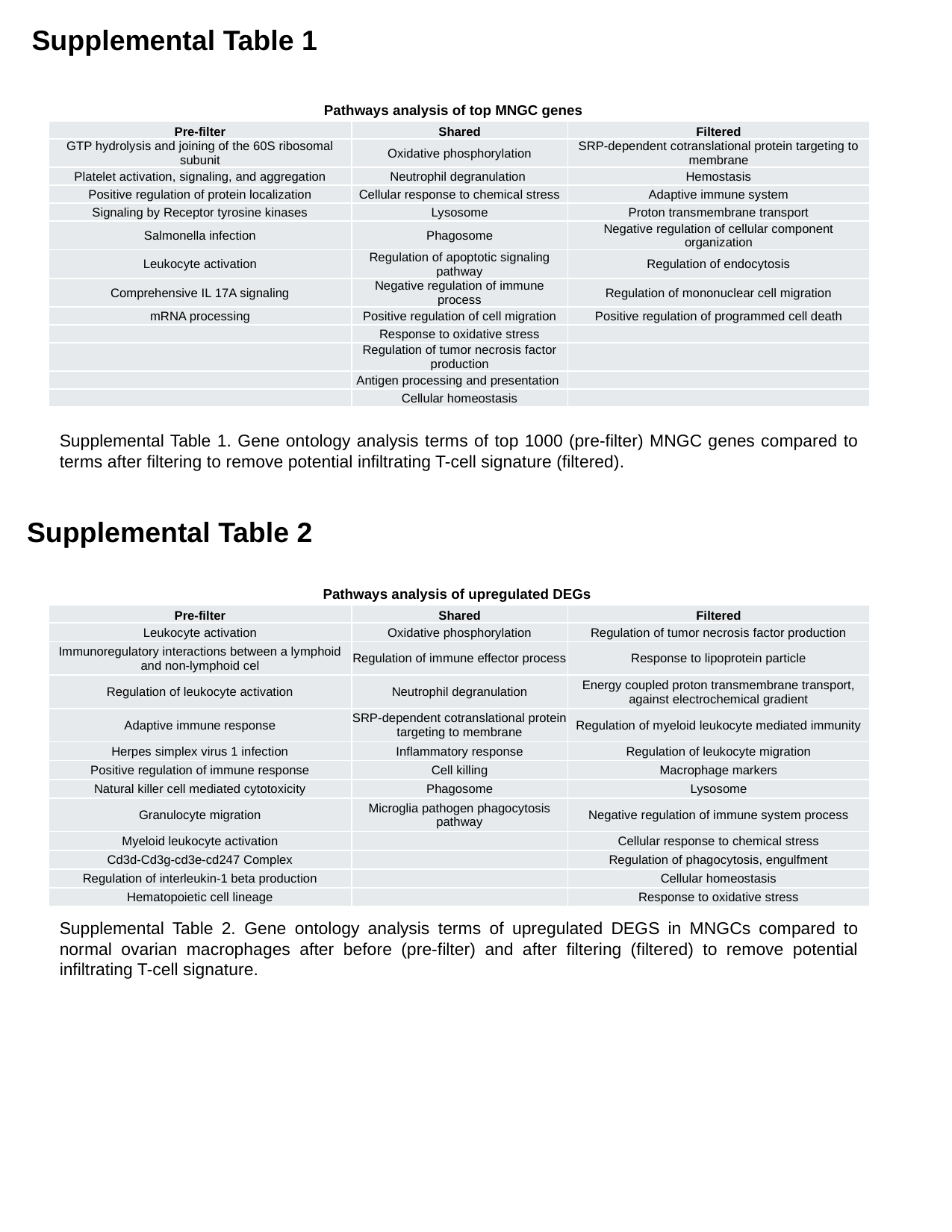

Supplemental Table 1
Pathways analysis of top MNGC genes
| Pre-filter | Shared | Filtered |
| --- | --- | --- |
| GTP hydrolysis and joining of the 60S ribosomal subunit | Oxidative phosphorylation | SRP-dependent cotranslational protein targeting to membrane |
| Platelet activation, signaling, and aggregation | Neutrophil degranulation | Hemostasis |
| Positive regulation of protein localization | Cellular response to chemical stress | Adaptive immune system |
| Signaling by Receptor tyrosine kinases | Lysosome | Proton transmembrane transport |
| Salmonella infection | Phagosome | Negative regulation of cellular component organization |
| Leukocyte activation | Regulation of apoptotic signaling pathway | Regulation of endocytosis |
| Comprehensive IL 17A signaling | Negative regulation of immune process | Regulation of mononuclear cell migration |
| mRNA processing | Positive regulation of cell migration | Positive regulation of programmed cell death |
| | Response to oxidative stress | |
| | Regulation of tumor necrosis factor production | |
| | Antigen processing and presentation | |
| | Cellular homeostasis | |
Supplemental Table 1. Gene ontology analysis terms of top 1000 (pre-filter) MNGC genes compared to terms after filtering to remove potential infiltrating T-cell signature (filtered).
Supplemental Table 2
Pathways analysis of upregulated DEGs
| Pre-filter | Shared | Filtered |
| --- | --- | --- |
| Leukocyte activation | Oxidative phosphorylation | Regulation of tumor necrosis factor production |
| Immunoregulatory interactions between a lymphoid and non-lymphoid cel | Regulation of immune effector process | Response to lipoprotein particle |
| Regulation of leukocyte activation | Neutrophil degranulation | Energy coupled proton transmembrane transport, against electrochemical gradient |
| Adaptive immune response | SRP-dependent cotranslational protein targeting to membrane | Regulation of myeloid leukocyte mediated immunity |
| Herpes simplex virus 1 infection | Inflammatory response | Regulation of leukocyte migration |
| Positive regulation of immune response | Cell killing | Macrophage markers |
| Natural killer cell mediated cytotoxicity | Phagosome | Lysosome |
| Granulocyte migration | Microglia pathogen phagocytosis pathway | Negative regulation of immune system process |
| Myeloid leukocyte activation | | Cellular response to chemical stress |
| Cd3d-Cd3g-cd3e-cd247 Complex | | Regulation of phagocytosis, engulfment |
| Regulation of interleukin-1 beta production | | Cellular homeostasis |
| Hematopoietic cell lineage | | Response to oxidative stress |
Supplemental Table 2. Gene ontology analysis terms of upregulated DEGS in MNGCs compared to normal ovarian macrophages after before (pre-filter) and after filtering (filtered) to remove potential infiltrating T-cell signature.

### Slide 4
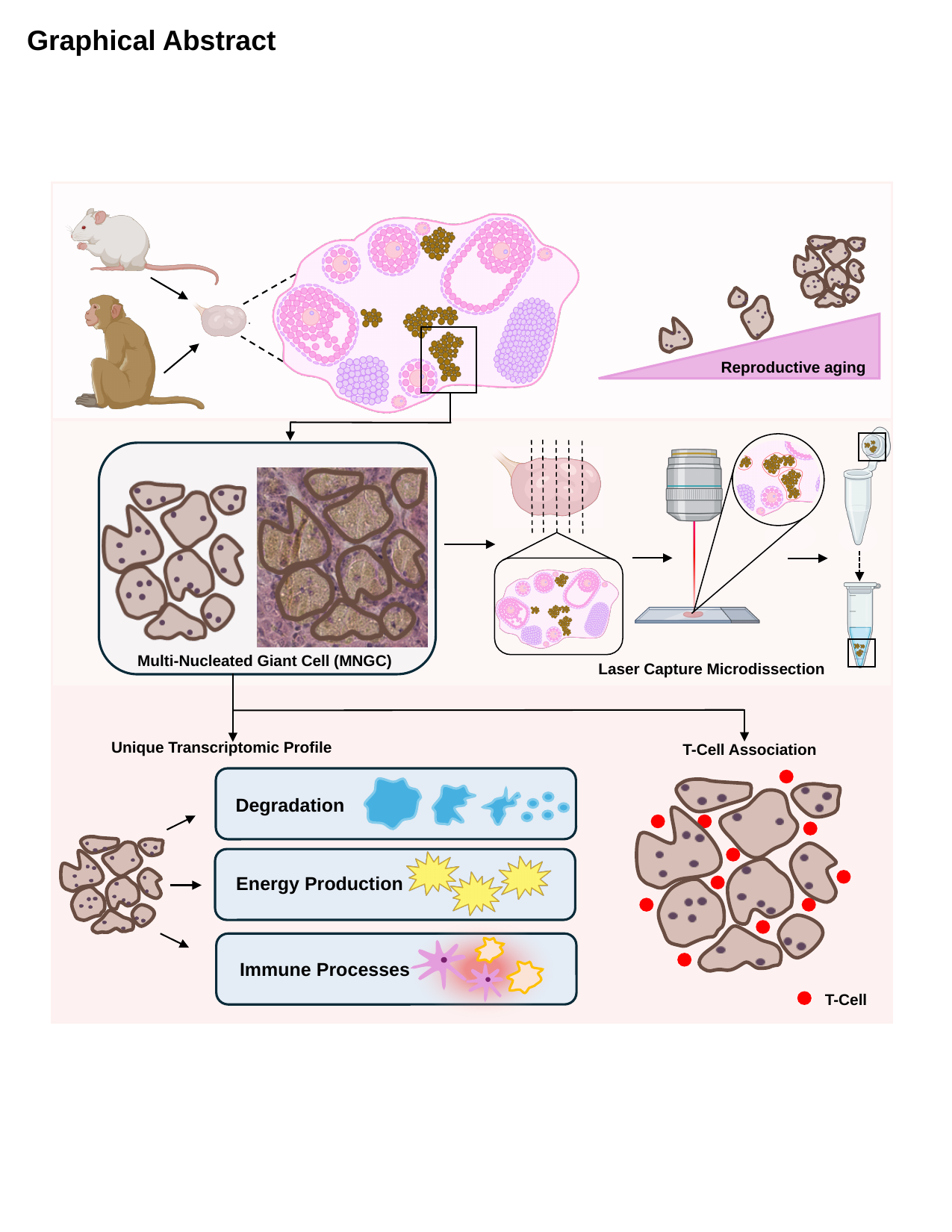

Graphical Abstract
Reproductive aging
Multi-Nucleated Giant Cell (MNGC)
Laser Capture Microdissection
Unique Transcriptomic Profile
T-Cell Association
Degradation
Energy Production
Immune Processes
T-Cell
